# The underlying mechanisms of improved balance after one and ten sessions of balance training in older adults

**DOI:** 10.1101/2020.10.01.322313

**Authors:** Leila Alizadehsaravi, Ruud A.J. Koster, Wouter Muijres, Huub Maas, Sjoerd M. Bruijn, Jaap H. van Dieën

## Abstract

Training improves balance control in older adults, but the time course and neural mechanisms underlying these improvements are unclear. We studied balance robustness and performance, H-reflex gains, paired reflex depression, and co-contraction duration in ankle muscles after one and ten training sessions in 22 older adults (+65yrs). Mediolateral balance robustness, time to balance loss in unipedal standing on a platform with decreasing rotational stiffness, improved (33%) after one session, with no further improvement after ten sessions. Balance performance, absolute mediolateral center of mass velocity, improved (18.75%) after one session in perturbed unipedal standing and (18.18%) after ten sessions in unperturbed unipedal standing. Co-contraction duration of soleus/tibialis anterior increased (16%) after ten sessions. H-reflex gain and paired reflex depression excitability did not change. H-reflex gains were lower, and soleus/tibialis anterior co-contraction duration was higher in participants with more robust balance after ten sessions, and co-contraction duration was higher in participants with better balance performance at several time-points. Changes in robustness and performance were uncorrelated with changes in co-contraction duration, H-reflex gain, or paired reflex depression. In older adults, balance robustness improved over a single session, while performance improved gradually over multiple sessions. Changes in co-contraction and excitability of ankle muscles were not exclusive causes of improved balance.

**Highlights:**

- Balance robustness and balance performance in perturbed unipedal standing was improved after one balance training session, with no further improvement after ten sessions.
- Balance performance in unperturbed unipedal standing was improved after ten sessions
- H-reflex and paired reflex depression did not change after training in unipedal or bipedal standing.
- Co-contraction duration of antagonistic ankle muscles increased after ten sessions in perturbed and unperturbed unipedal standing.
- Changes in co-contraction duration and excitability of ankle muscles were not exclusive causes of improved balance.

## 1. Introduction

Balance control is essential to avoid falls during daily-life activities. Impaired balance control due to aging results in falls, injuries, and loss of independence in older adults (World Health Organization, 2007). To resolve this issue, it is important to understand how balance control works and when and how it improves as a result of training. Balancing requires the central nervous system to act rapidly and accurately on an array of sensory inputs (Chiba et al., 2016), consisting of visual, vestibular, and tactile information, as well as proprioceptive sensory feedback (Inglis et al., 1994). Balance training leads to improved balance performance in older adults (Muehlbauer et al., 2015), observed as a reduction in mediolateral center of mass velocity during unipedal stance (Raymakers et al., 2005). However, the question of how balance training induces changes in neuromuscular control remains unanswered. Hence, it is important to investigate the relation between improved balance control in older adults with changes in neural mechanisms at central and/or peripheral nervous system components.

Changes in balance control with training appear to occur at short-time scales, with substantial improvements after a single trial and over a single session (Liu et al., 2017; Rieger et al., 2020; van Dieën et al., 2015). Previously a rapid improvement of balance control in young adults after one session of balance training has been shown (van Dieën et al., 2015), while results of short-term training in older adults were inconsistent (Berghuis et al., 2015; Coats et al., 2014) and most studies have focused on training over several sessions spread over multiple weeks (Muehlbauer et al., 2015). Since training effects were measured mainly before and after the entire training period only, the difference between a single session and several sessions of balance training in older adults is unclear.

In addition, most studies assess balance control with measures that capture balance performance, which quantifies how good people are at minimizing disturbances from an equilibrium position (often in a non-challenging condition like bipedal stance or unipedal stance on a rigid surface), for instance, by measuring postural sway, in which lower values indicate better performance (Cadore et al., 2014; Leiros-Rodríguez & García-Soidan, 2014; Martinez-Amat et al., 2013; Yu et al., 2014). There are two problems with this. Firstly, subjects may choose not to minimize their sway, as higher sway values may be unproblematic and require less energy (Houdijk et al., 2015). Secondly, even if subjects choose to minimize their sway, balance performance does not reflect the capability to avoid balance loss when challenged, i.e., balance robustness, which quantifies the largest perturbation that can be resisted. Robustness has received limited attention in training literature, and if it is measured, it is mostly done in a dichotomous way (ability to perform a task, e.g., stand on one leg with eyes closed for 10 s, or not) (Pasma et al., 2014). For practical purposes, improved robustness may be more important than improved performance. While improved balance performance may not necessarily prevent falls, it may indicate improvements in balance control. Hence, we here chose to study the effects of training on both these aspects of balance control.

Age-related degenerative processes in the sensory and motor systems induce a shift from reliance on feedback control to reliance on feedforward strategies, reflected in increased intensity of co-contraction of antagonistic ankle muscles (Benjuya et al., 2004). In a simulation study, such increased antagonistic co-contraction was shown to compensate for impaired sensory feedback (Koelewijn & Van Den Bogert, 2020). Also young adults, increase both the intensity and duration of antagonistic co-contraction when confronted with a challenging balance task, (Alizadehsaravi et al., 2020; Gebel et al., 2019). Moreover, higher antagonistic co-contraction duration and magnitude, were found in older adults with poor balance control compared to young adults with better balance control (Alizadehsaravi et al., 2020; Nagai et al., 2011). Balance training can potentially reduce magnitude and duration of antagonistic co-contraction (Schinkel-Ivy & Duncan, 2018). We note here that many different methods have been used to assess co-contraction in the literature. In the studies cited, the index of co-contraction reflected either the magnitude of co-contraction (Gebel et al., 2019) or its magnitude and duration combined (Alizadehsaravi et al., 2020; Nagai et al., 2011; Schinkel-Ivy & Duncan, 2018).

Alterations of the H-reflex indicate an adjusted motoneuron output after processing of la afferent input at the spinal cord (Payton, 1978). With age, postural modulation of H-reflexes is reduced (Alizadehsaravi et al., 2020; Earles et al., 2000; Koceja et al., 1995; Koceja & Mynark, 2000) and this may be functionally related to a declined balance performance in older adults (Mynark & Koceja, 2001). Balance training in young adults has been reported to decrease the soleus (SOL) H-reflex (Keller et al., 2012; Taube, Gruber, et al., 2007; Taube, Kullmann, et al., 2007; Trimble & Koceja, 2001). While both young and older adults are capable of down-training the SOL H-reflex (Mynark & Koceja, 2002), it is unclear whether balance training also causes such down-regulation of the H-reflex in older adults. Unfortunately, only a few studies have addressed the effect of training on the H-reflex in older adults. Scaglioni et al. showed no changes in the H-reflex after 16 weeks of strength training in older adults (Scaglioni et al., 2002), Ruffieux et al. found no effects of training on H-reflex after five weeks (Ruffieux et al., 2017), and Lauber et al. showed an enhanced H-reflex after 12 weeks of alpine skiing (Lauber et al., 2011). Decreases in the H-reflex are thought to reflect a reduced effect of spinal feedback circuitry on motor control, coinciding with increased supraspinal control (Baudry et al., 2015; Taube, Gruber, et al., 2007). However, supraspinal mechanisms also affect the excitability of the alpha motoneuron pool and, therefore the H-reflex gain. This hampers the interpretation of the H-reflex. Therefore, measurements of paired reflex depression were added in the current study to provide an insight into peripherally induced inhibition which would more exclusively reflect changes in peripherally induced presynaptic inhibition (Jeon et al., 2007; Sefton et al., 2007; Trimble et al., 2000). The second H-reflex in paired reflex depression measurements is assumed to be influenced by the synchronous activation of the spindle’s afferents during the first H-reflex. Using paired reflex depression, the influence of primary spindle afferent feedback and therefore, activation history of the Ia afferents on the motoneuron pool output can be studied (Trimble et al., 2000). Among middle-aged adults (^~^44 years), subjects with long-term Tai Chi practice showed better balance performance and, despite a similar H-reflex, a larger paired reflex depression (Guan & Koceja, 2011). These authors assumed that a reduced second H-reflex avoids overcorrection and prevents unwanted oscillations. Hence, increased paired reflex depression might be expected as a result of balance training.

The aims of the present study were twofold; first, we aimed to assess the functional benefits of one session and ten sessions of balance training in older adults. To do so, we assessed changes in balance robustness (as the duration that participants were able to keep their body balanced while surface stiffness was decreased) and balance performance (measured as the mean absolute value of the mediolateral center of mass velocity during unipedal balancing). Second, we aimed to explore the associations between the changes in balance robustness and balance performance with co-contraction duration, H-reflex gain, and paired reflex depression after one session and ten sessions of training. We hypothesized that balance robustness and performance would be improved slightly after one session, and significantly after ten sessions of training and that such improvements would be accompanied by changes, such as decreased antagonistic co-contraction duration, lower H-reflex gains, and stronger paired reflex depression.

## 2. Methods

### 1.1 Participants

Twenty-two healthy older adults (age: 72.6 ± 4.2 years, length: 1.71 ± 0.09 m, weight: 75.6 ± 13.3 kg; mean ± SD, 11 females and 11 males) participated in this study. This is comparable to similar studies (Bisson et al., 2007; Nagy et al., 2007) and in accordance with a required sample size of twenty-two based on power analysis for an F test of a within factor repeated measure, assuming an effect size of 0.44 (Muehlbauer et al., 2015) and correlation among repeated measures of 0.6 (β = 0.8, G*power 3.1.9.2, Düsseldorf, Germany). To ensure participant safety and data reliability, exclusion criteria included: an inability to stand and walk for 3 minutes without walking aid; cognitive impairments (MMSE <24); depression (GFS > 5); obesity (BMI > 30); orthopedic, neurological, and cardiovascular disease; use of medication that affects balance; and severe auditory & visual impairments. To prevent ceiling effects in balance robustness and performance and limited training gains, participants practicing sports that explicitly include balance components (e.g., Yoga, Pilates) were excluded as well (Kiers et al., 2013). To prevent obscuring any training effects, participants were asked to keep their normal activity levels in their daily life throughout the experiment. All participants provided written informed consent prior to participation, and the experimental procedures were approved by the ethical review board of the Faculty of Behaviour and Movement Sciences, Vrije Universiteit Amsterdam (VCWE-2018-171).

### 1.2 Experimental procedures

The protocol included an initial measurement session to determine baseline state (Pre), a measurement after one 30-minutes session of balance training (Post1), and after ten 45-minutes sessions (Post2). The protocol was concluded with a retention assessment (Ret) two weeks after the last training session. The Pre-measurements, the 30-min training session, and the Post1 measurements were performed on the same day. The measurements consisted of blocks of tasks and tests in the following order: familiarization, assessment of balance robustness, baseline electromyography measurement (EMG; only at Pre and Post2), reflex assessment, and a series of (perturbed and unperturbed) unipedal balance performance tests. During the reflex assessments and the series of unipedal balance performance tests, kinematic and EMG data were recorded. The retention measurement consisted solely of the assessment of balance robustness (see Figure 1 for an overview).

**Figure 1:**
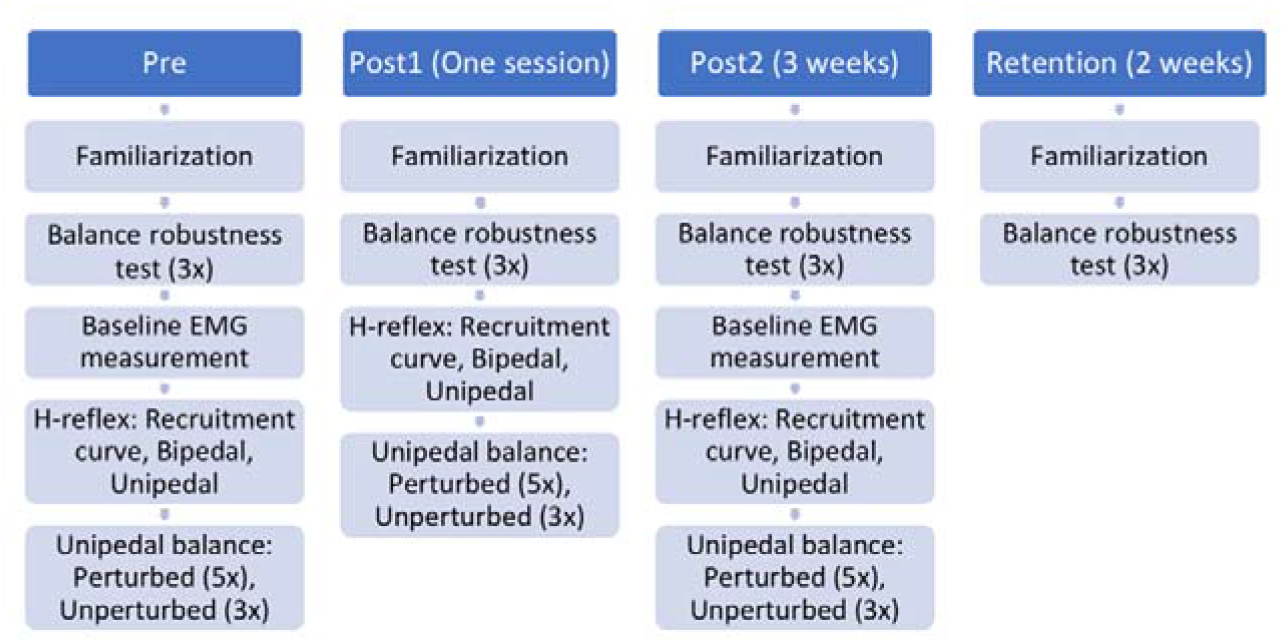
Diagram illustrating the experimental procedures.

### 1.3 Instrumentation and data acquisition

For all unipedal tasks, a custom-made balance platform controlled by a robot (HapticMaster, Motek, Amsterdam, the Netherlands) was used. This platform can rotate 17.5° to either direction in the frontal plane. The rotation of the platform can be controlled by the robot, either simulating a tunable stiffness, expressed in Nm/rad, or applying orientation control. During tasks where the platform simulated a stiffness, the rotational stiffness of the platform was normalized to percentage of mgh (body weight multiplied by center of mass height) of each participant, to factor out differences in participant height and mass. Stiffness approaching zero allows the platform to freely rotate in the frontal plane, similar to standing on a wobble board with a small radius of curvature, while a high stiffness resembles standing on a rigid surface. For safety reasons, the balance platform was equipped with bars in front and on both sides of the participant, and there was ample space to step off the rotating part of the platform (Figure 2). Participants were instructed to keep their arms slightly abducted at the sides of their body and to freely use their arms to maintain their balance if needed. Also, they were allowed to grab or touch the bars in case of balance loss.

**Figure 2:**
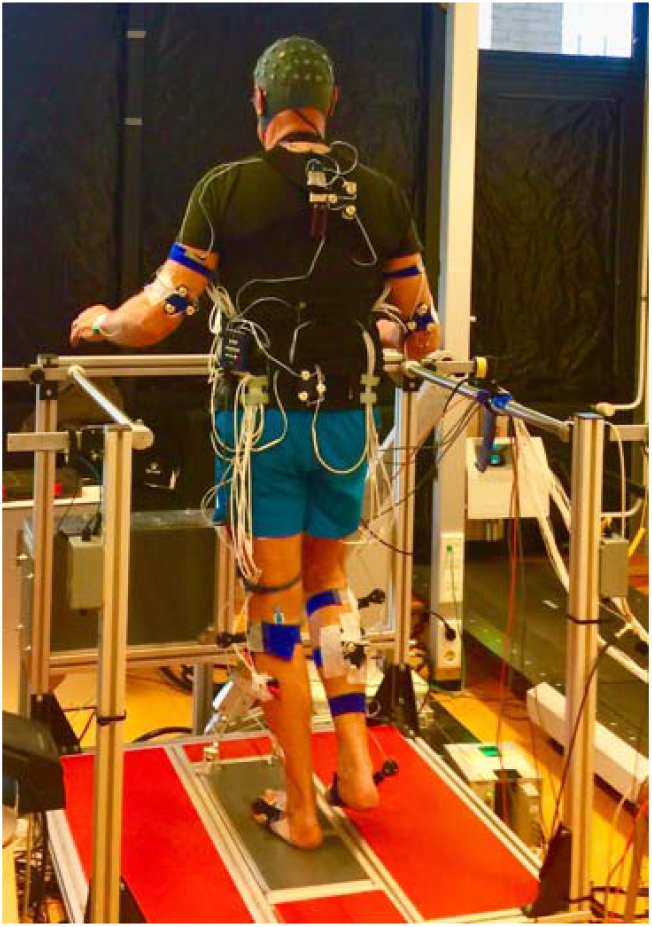
Participant in unipedal stance on the robot-controlled balance platform. (This article is a part of a larger study; EEG data will be reported later).

Surface EMG data were collected from three muscles on the preferred stance leg: m. tibialis anterior (TA), m. peroneus longus (PL), and m. soleus (SOL). PL was selected for its role in mediolateral ankle movement (eversion). TA and SOL have effects on anteroposterior balance control, but due to the orientation of the talocalcaneal joint axis, TA and SOL both also have a mediolateral (eversion/inversion) component. The TA is even considered to be the main agonists for inversion. In addition, the TA and SOL muscles may facilitate stronger or faster responses of for example the PL muscle, which has an effect in anteroposterior control as well. Bipolar electrodes were placed in accordance with the SENIAM recommendations (Hermens et al., 2000). The EMG signals were sampled at 2000 Hz and amplified using a 16-channel TMSi Porti system (TMSi, Twente, The Netherlands). The baseline EMG was measured during unipedal stance on a rigid surface. The preferred stance leg was reported by the participant prior to the experiment and confirmed by the experimenter by asking the participant to kick an imaginary soccer ball. The supporting leg was considered the preferred stance leg.

Kinematic data were obtained from 8 active marker clusters, containing three markers each, placed on the posterior surface of the thorax (1), pelvis (1), arms (2), calves (2), and feet (2). The trajectories of these clusters were tracked by one Optotrak camera array (Northern Digital, Waterloo, Canada). An anatomical model of the participants was constructed from the 8 cluster-markers by relating the cluster location and orientation on each segment to a minimum of three anatomical bony landmarks (see supplementary material 1) for that segment, using a four-marker probe, while standing in upright position. Based on the assumptions of rigid and connected body segments, as well as the cluster trajectories, a kinematic model was formed (Cappozzo et al., 1995).

#### 1.3.1 Familiarization

At Pre and Post2 time-points, participants were familiarized with standing on the platform on their preferred leg in two trials. In the first familiarization trial, the platform imposed ten 8° rotational perturbations at a rate of 16°/s in random direction and returned to horizontal state, every 3 s, to familiarize the subjects with perturbed unipedal balancing. In the second familiarization trial, the platform was set at a stiffness of 100% mgh for 30 s, while participants had to remain balanced.

#### 1.3.2 Balance robustness test

Unipedal balance robustness was assessed using the balance platform. After familiarization and rest, the participants had to stand on their preferred leg until balance loss occurred, while the stiffness of the platform decreased stepwise every 5 s, asymptotically approximating 0 Nm/rad at the maximum trial duration of 100 s (see EQ. 1, Figure 3). The time an individual could stay balanced, without grabbing the bar or putting down the other foot, was used to assess balance robustness. This was repeated three times, with ample rest (2-5 minutes) in between, and results were averaged over three trials.

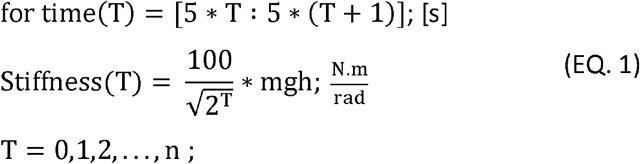

**Figure 3:**
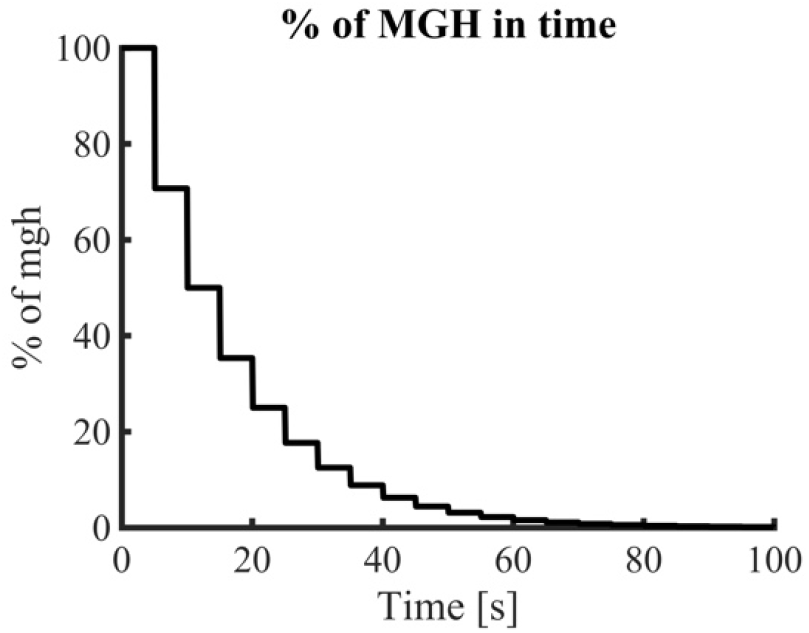
The duration and corresponding stiffness of the test for robustness.

#### 1.3.3 Baseline EMG measurement

The EMG baseline was measured during unipedal stance on a rigid surface for 60 s. This recording served as baseline measurement for co-contraction duration. For Pre and Post1 the same baseline measurement was used, as they were recorded on the same day.

#### 1.3.4 Balance performance tests

To assess balance performance two unipedal balance trials were performed on the robot-controlled platform: an unperturbed and a perturbed task. In the unperturbed task, the stiffness of the platform was set at a constant value. To normalize task difficulty to individual skill level, the stiffness was set at 1.3 times the value at which balance loss occurred during the assessment of balance robustness in the Pre-measurement. This task was repeated three times with two minutes rest between trials. In the perturbed task, twelve perturbations were imposed by the platform in the form of mono-phasic sinusoidal rotations either in medial or lateral direction (amplitude of 8°, angular speed of 16°/s). The perturbation direction was randomized, and the inter-perturbation duration was randomly selected between 3-5 s to prevent anticipatory behaviour. This task was performed five times with two minutes rest in between trials.

#### 1.3.5 Reflex assessment

To elicit the H-reflex in the SOL, the tibial nerve was stimulated using an electrical stimulator (Digitimer, DS7A UK). A large diameter anode, roughly 6 × 9 cm constructed of aluminum foil and conducting gel, was fixed on the patella of the standing leg (Zehr, 2002). The cathode was placed over the tibial nerve in the popliteal fossa of the same leg. The optimal cathode position was determined in each subject by probing the popliteal fossa and delivering 5-10 mA stimulations to find the location that resulted in the largest SOL H-reflex amplitude ^~^25 ms after stimulation.

Assessment of the H-reflex consisted of three parts: determining the recruitment curve to find H_max_ and M_max_, measuring the H-reflex and paired reflex depression in bipedal stance (bipedal reflex trial), and measuring H-reflex and paired reflex depression in unipedal stance (unipedal reflex trial), with the intensity of the stimulator set at H_max_ for the latter two parts. To obtain the recruitment curve, participants were subjected to low-amplitude (^~^5 to ^~^120 mA) electrical stimuli. Participants were instructed to stand still bipedally, with the feet placed at shoulder width, arms besides their body, and to focus their vision on a target in front of them. Subsequently, 1 ms single square pulses with a minimum 4 s inter-stimulus duration were delivered to the tibial nerve at increasing amplitudes to elicit H-reflexes in the SOL while EMG data was recorded. H_max_ is the maximum peak-to-peak amplitude of the SOL EMG, between 25 and 50 ms post stimulation, and M_max_ is the maximum peak-to-peak amplitude of SOL EMG between 0 and 25 ms post stimulation.

Subsequently, H-reflex and paired reflex depression were assessed in two stance conditions, (1) stable bipedal stance (bipedal reflex trial), and (2) unipedal stance (unipedal reflex trial) on the balance platform with the stiffness set at 100% mgh (Jeon et al., 2007). In both conditions, participants were subjected to ten double-pulse stimulations of the tibial nerve. Here, inter-pulse duration was 100 ms, inter-train duration was randomized between 4-8 s, and stimulation intensity was set to the level that previously elicited the H_max_. The first H per double-pulse was considered the H-reflex.

#### 1.3.6 Balance training

In the first session, the participants were trained individually. The nine sessions of the 3-week training program took place in a group setting (6-8 participants). The training program was designed based on previous studies that reported improved balance and reduced fall-risk (Lesinski et al., 2015; Sherrington et al., 2017). All training sessions were supervised by a physical therapist who ensured that the sessions remained safe, yet sufficiently challenging, for all participants. The difficulty of the exercise was manipulated by: reducing support (e.g. hand support, two-legged stance, unipedal stance), using unstable objects with varying degrees of freedom and stability, adding motor and cognitive tasks (e.g., catching a ball or passing it in changing directions), and reducing sensory information (e.g., visual fixation or eyes closed). Each session started with a short warm-up. Solely standing balance exercises focusing on unipedal stance were included in the training program (Giboin et al., 2015), (see supplementary material 2). Group training sessions were 15 minutes longer than individual training sessions. Extra time was required to switch the devices between the training partners in the exercises with equipment.

## 2. Data analysis

### 2.1 Balance robustness

The duration the participant maintained balance, averaged over three trials, served to assess the individual’s balance robustness.

### 2.2 Balance performance

The trajectory of the center of mass (CoM) was estimated from the full body kinematic model (Kingma et al., 1996). Balance performance was expressed as the mean absolute center of mass velocity in the mediolateral direction (vCoM).

### 2.3 Co-contraction duration (CCD)

Co-contraction is the concurrent activation of two muscles. It can be expressed as the duration, magnitude, or both duration and magnitude of concurrent activation. In our study co-contraction duration was derived from two muscle pairs: SOL/TA, and TA/PL. The raw EMG data were high-pass (35 Hz, bidirectional, 2^nd^ order Butterworth) and notch filtered (50 Hz and its harmonics up to the Nyquist frequency, 1 Hz bandwidth, bidirectional, 1^st^ order Butterworth). Subsequently, the filtered data were rectified using the Hilbert transform and low-pass filtered (40 Hz, bidirectional, 2^nd^ order Butterworth). Finally, we determined the percentage of data points during the perturbed and unperturbed tasks at which both muscles in a pair exceeded the mean muscle activity of baseline unipedal stance (baseline EMG measurement). The baseline measurement was used to set an arbitrary but consistent threshold for the index of co-contraction, which reflects the duration during which both muscles are substantially active (i.e., more active than in quiet unipedal standing).

Since for Pre and Post1 time-points the measurements were performed on the same day, the same unipedal baseline trial was used as a reference for these two time-points.

### 2.4 H-reflexes and Paired Reflex Depression (PRD)

For H-reflex gain and paired reflex depression the raw EMG of the SOL was high-pass filtered (10 Hz, bidirectional, 2^nd^ order Butterworth). The H-reflex gain (EQ. 2) was calculated as the mean, over all pulse trains.

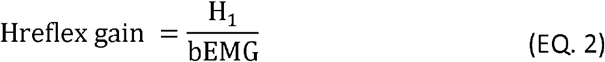

where H1 was the maximum peak-to-peak amplitude ^~^25 ms after the first stimulus of the paired-pulse train and bEMG was the root-mean-square value of the EMG activity over the 100 ms prior to the pulse train. paired reflex depression was quantified as the mean relative depression of the second H-reflex relative to the first one (EQ. 3).

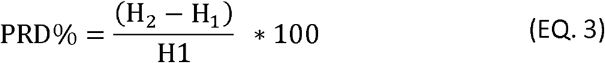

### 2.5 Statistics

A one-way repeated measures ANOVA was used to test the main effect of time-point (Pre, Post1, Post2, Retention) on balance robustness. Post-hoc comparisons (paired sample t-tests) were performed to investigate the effect of one session of training (Pre vs. Post1), long-term training (Pre vs. Post2), and retention (Pre vs. Retention). In addition, Post1-Post2 and Post2-Retention were compared to obtain insight into the changes over the short- and long-term and in retention.

Two-way repeated-measures ANOVAs were used to identify main effects of time-point (Pre, Post1, Post2) and condition (perturbed/unperturbed or bipedal/unipedal reflex trials) on vCoM, co-contraction duration, H-reflex gain, and paired reflex depression. When the assumption of sphericity was violated, the Greenhouse-Geisser method was used. Post-hoc analyses (paired samples t-test) were performed to investigate the effect of one session (Pre vs. Post1) and ten sessions (Pre vs. Post2) of training when a main effect of Time-point or an interaction of Time-point × Condition was observed. For all post-hoc analyses, Holms’ correction for multiple comparisons was applied.

Balance performance and the response to training are heterogeneous in older adults (Muehlbauer et al., 2015). Therefore, cross-sectional and longitudinal correlation analyses were performed to gain more insight into which (changes in) co-contraction, H-reflexes, and paired reflex depression were related to (changes in) balance robustness and balance performance. As cross-sectional analyses, the correlations between balance robustness (duration) and co-contraction duration (averaged over perturbed and unperturbed trials) for both muscle pairs, H-reflex, and paired reflex depression were calculated. In view of a positive correlation between co-contraction in perturbed and unperturbed trials, we averaged the results and calculated the correlation between co-contraction duration (perturbed + unperturbed) and robustness.

Moreover, the correlations between balance performance (vCoM) and the co-contraction duration for both muscle pairs in perturbed and unperturbed trials and between balance performance (vCoM) and H-reflex gains and paired reflex depression during unipedal reflex trial and bipedal reflex trial were calculated for the three time-points. For longitudinal analyses, the correlations between changes in the same parameters after one session and ten sessions of training were calculated. In view of outliers, Spearman’s correlation (r) coefficients were calculated. In all statistical analyses, α=0.05 was used.

Only in balance robustness test, all participants were included in the analysis. For all other analyses twenty-one participants were included because one participant was not able to fully perform the balance performance trials.

## 3. Results

### 3.1 Balance robustness

Balance robustness (duration of balancing) increased as a result of balance training (F_1.955,41.060_= 10.637, p < 0.001). The mean duration of balancing increased after one session of training (t = 3.325, p = 0.006, Figure 4). While the duration remained unchanged between Post1-Post2 and Post2-Retention (t = − 1.257, p = 0.427; t = − 0.57, p = 0.571, respectively; Figure 4), ten sessions of training and retention showed higher robustness than Pre time-point (t = − 4.582, p <0.001; t = − 5.151, p <0.001, respectively; Figure 4). Overall, these results indicate a rapid improvement in balance robustness after only one session of training, with no further improvement after the subsequent nine training sessions.

**Figure 4:**
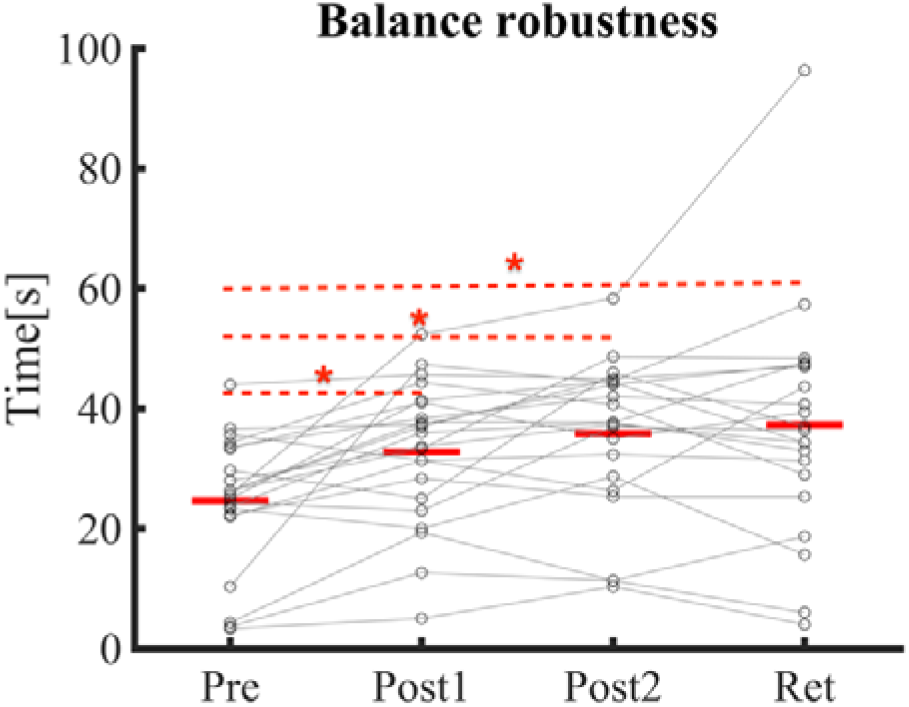
Balance robustness at different time-points, expressed as the duration of maintaining balance under gradually decreasing surface stiffness. The red asterisk (*) indicate statistical significance.

### 3.2 Balance performance

Balance training led to an increase in balance performance (decreased vCoM; Figure 5; Time-point effect, F_1.533,30.655_= 10.598, p < 0.001). Participants showed larger vCoM in perturbed compared to unperturbed standing (Figure 5.a & 5.b, Condition effect, F_1,20_ = 58.285, p <0.001). Additionally, there was a significant interaction of time-point and condition on vCoM (F_2,40_= 5.242, p = 0.01). Post-hoc analysis showed that one session of training decreased vCoM in the perturbed condition but did not change vCoM in the unperturbed condition (t = 3.35, p = 0.011 and t = 1.193, p = 0.715, respectively). On the other hand, ten sessions of training changed vCoM significantly in both perturbed and unperturbed condition (t = 5.206, p <0.001; t = 3.394, p = 0.011, respectively; Figure 5.a & 5.b), even though there were no significant changes in vCoM between Post1 and Post2 measurements in perturbed and unperturbed conditions (t = 1.439, p = 0.783; t = 1.718, p = 0.553, respectively; Figure 5.a & 5.b).

**Figure 5:**
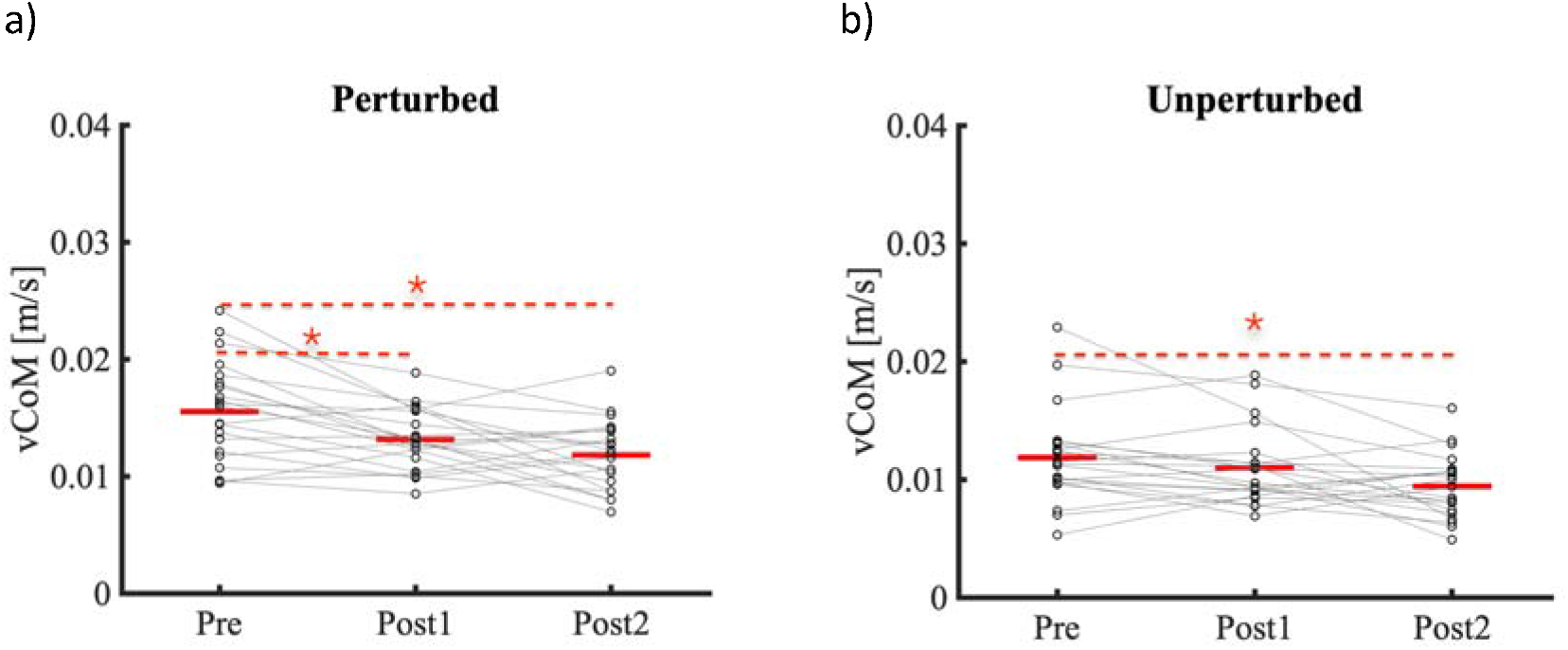
The mean absolute center of mass velocity in mediolateral direction at all three measured time-points. a) in the perturbed condition, b) in the unperturbed condition. The red lines indicate averages across subjects. The red asterisk (*) indicate statistical significance.

### 3.3 Neural mechansisms

#### 3.3.1 Duration of Co-contraction

In perturbed and unperturbed trials, co-contraction duration of SOL/TA muscle pair was affected by time-point (Table 1). Post-hoc comparison showed that the co-contraction duration was not changed after the one session but was increased after ten sessions of training (t = 1.623, p = 0.112; t = − 2.372, p = 0.045, respectively; Figure 6.a). No effects of Time-point and Condition, nor an interaction were observed for TA/PL (Table 1). Overall, our results showed no changes in SOL/TA co-contraction duration after one session of training but an increased SOL/TA co-contraction duration after ten sessions of training.

**Table 1:**
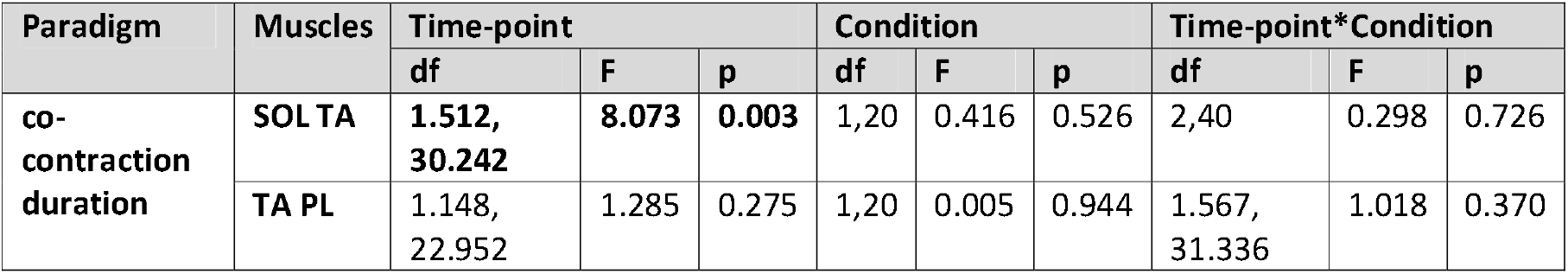
Results of repeated-measures ANOVA of the duration of co-contraction of two muscle pairs, in perturbed and unperturbed standing at three Time-points of Pre, Post1, and Post2. Bold numbers indicate a significant effect.

**Figure 6:**
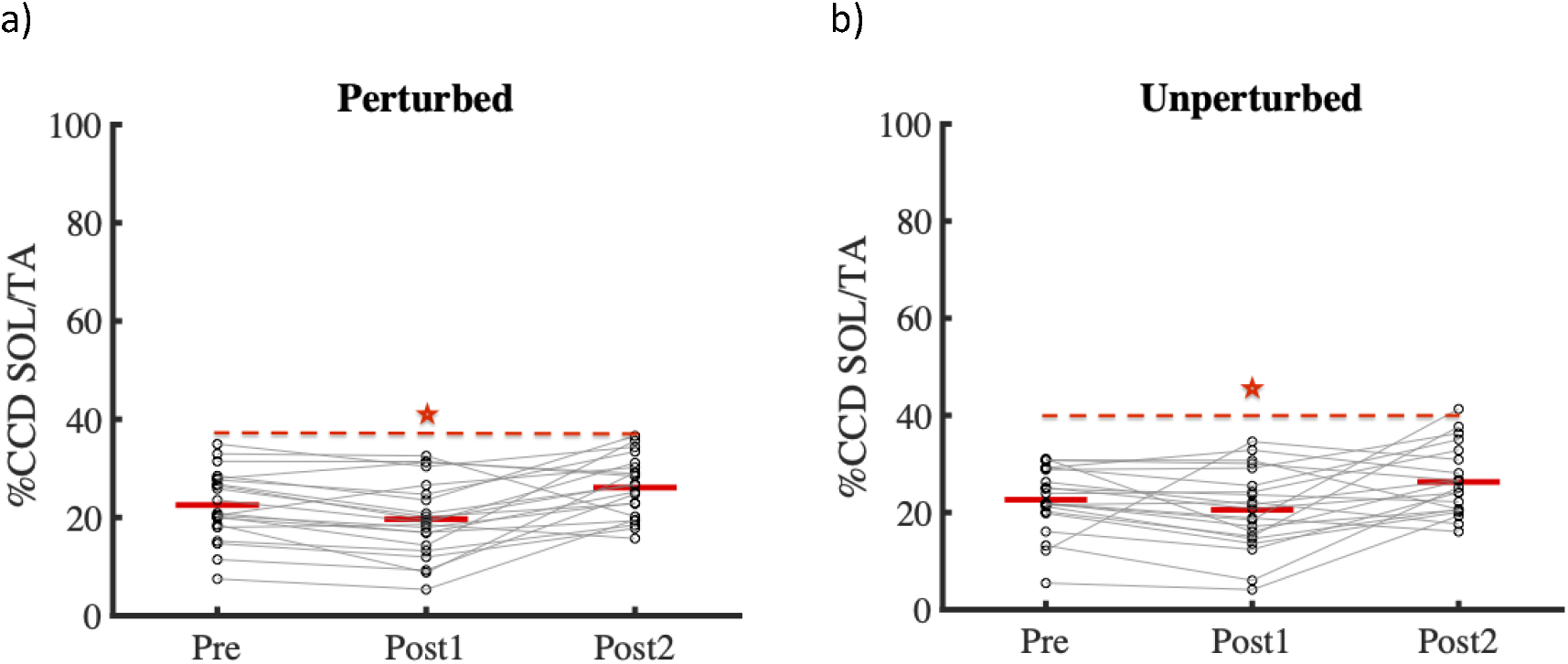
Co-contraction duration at three time-points in a) perturbed and b) unperturbed standing, for the muscle pairs SOL/TA. Circles and connecting lines represent individual results. The red lines indicate averages across subjects. The red asterisk (*) indicate statistical significance.

#### 3.3.2 Reflexes

There was no effect of Time-point, nor an interaction effect of Time-point x Condition, on H-reflex gains (F_1.567,31.344_= 0.467, p = 0.585, and F_2,40_ = 1.859, p = 0.169, respectively; Figure 7). H-reflex gains were significantly higher in bipedal compared to unipedal reflex trial (F_1,20_= 26.549, p <0.001). Similarly, there was no effect of Time-point, nor an interaction effect of Time-point x Condition, on paired reflex depression (F_2,40_= 1.043, p = 0.360, and F_2,40_= 0.204, p = 0.802, respectively; Figure 8), but paired reflex depression was stronger in bipedal compared to unipedal reflex trial (F_1,20_= 39.613, p <0.001). Overall, our results did not show any changes in the reflexes as a result of training.

**Figure 7:**
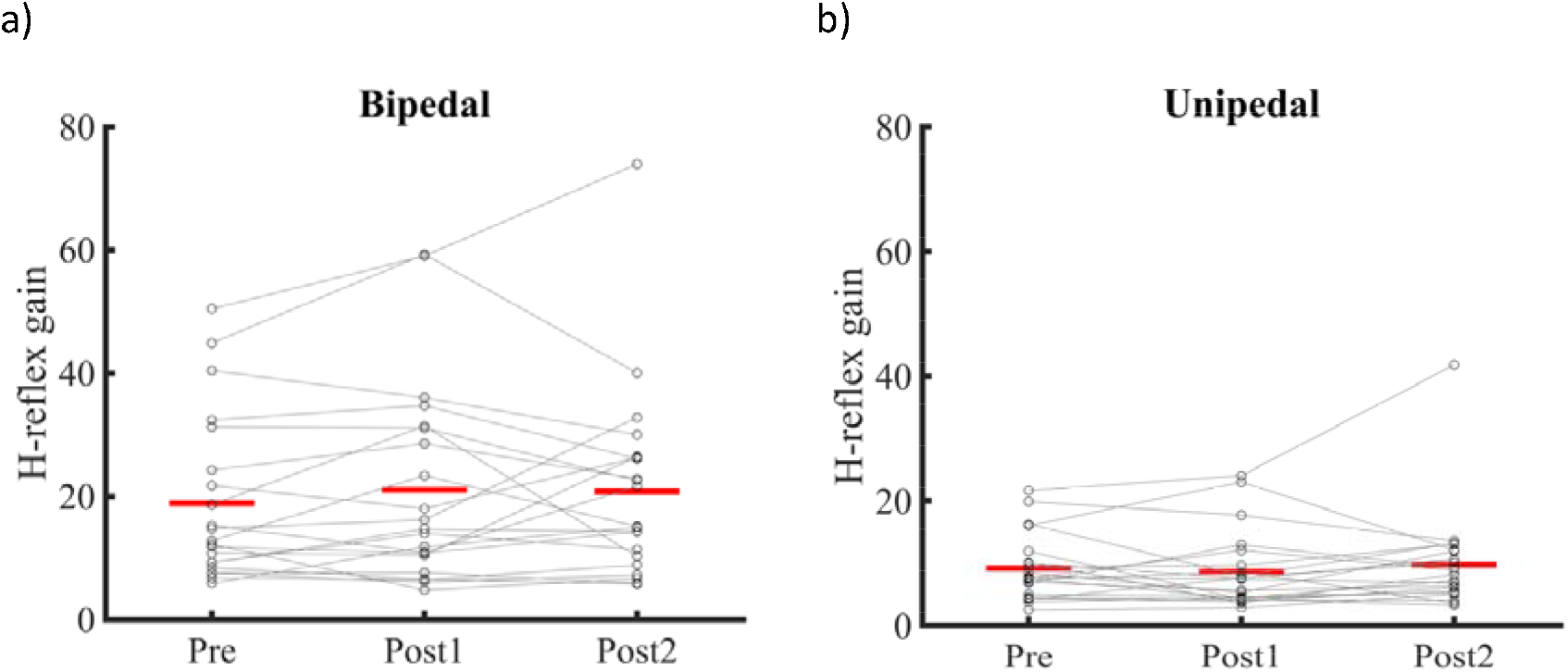
H-reflex gains at three time-points a) shows the reflex gain for the bipedal reflex trial b) shows the reflex gain for the unipedal reflex trial. Circles and connecting lines represent individual results. The red lines indicate averages across subjects.

**Figure 8:**
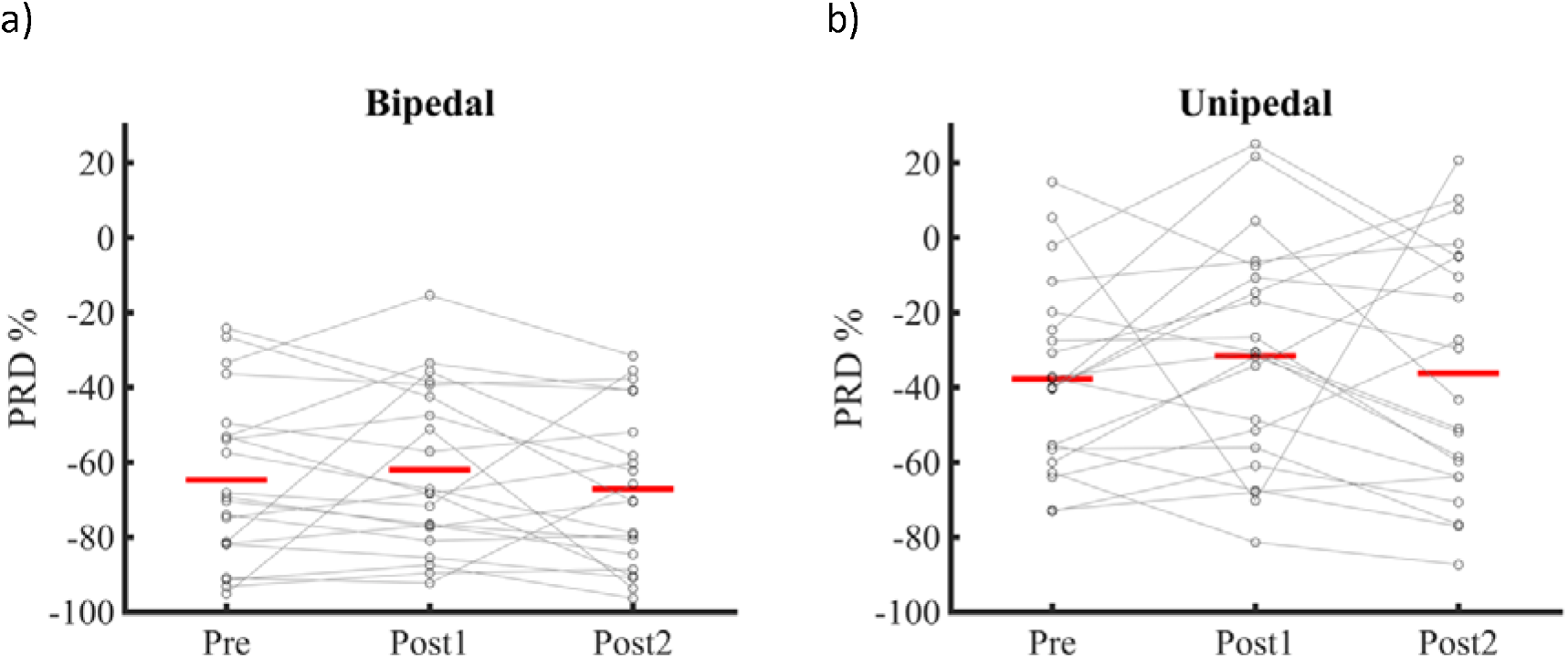
Paired reflex depression at three time-points. The paired reflex depression is displayed for a) the bipedal reflex trial, and b) unipedal reflex trial. Circles and connecting lines represent individual results. The red lines indicate averages across subjects.

#### 3.3.3 Associations of balance robustness with co-contraction and reflexes

All correlation results are shown in Tables 2 and 3. For co-contraction, the average of the perturbed and unperturbed SOL/TA co-contraction duration was positively correlated with balance robustness at time-point Post2 (r = 0.564, p = 0.007). No correlations were observed between changes after one session or ten sessions of training. For reflexes, H-reflex gains in unipedal stance were negatively correlated with balance robustness (duration) at time-point Post2 (r = −0.585, p = 0.005). No correlations were observed between changes after one session or ten sessions of training.

**Table 2:**
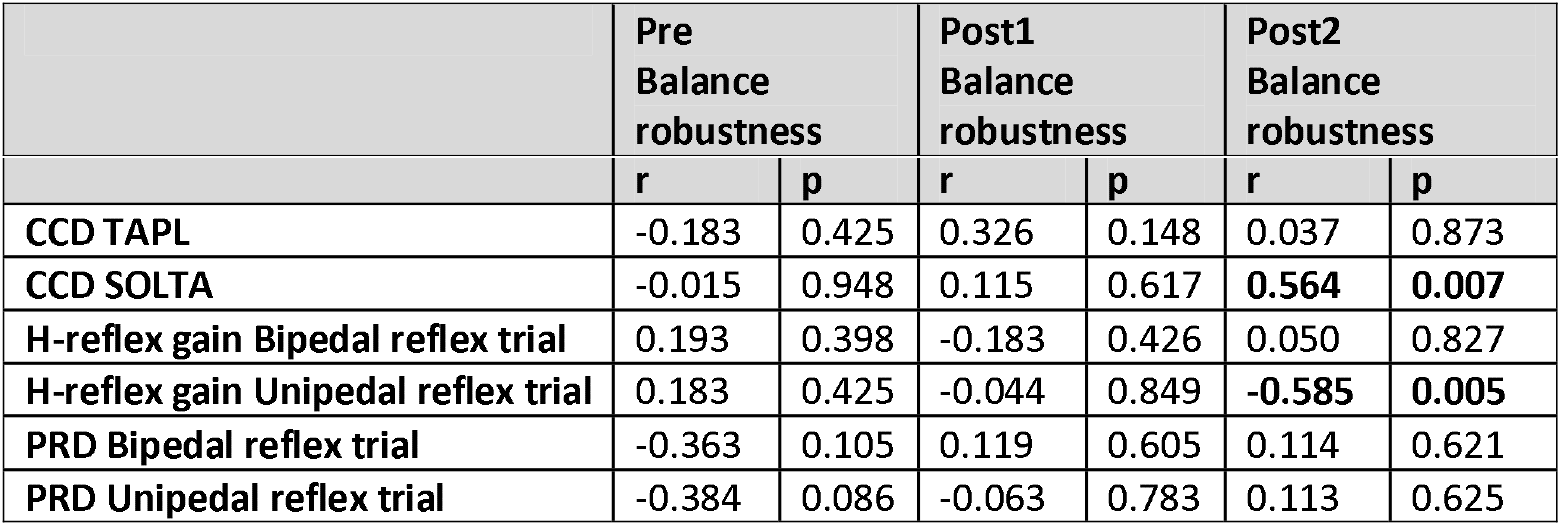
Results of the correlational analysis between co-contraction (averaged over perturbed and unperturbed trials), reflexes in bipedal and unipedal reflex trials with balance robustness (duration) at each Time-point. Bold numbers indicate a significant effect.

**Table 3:**
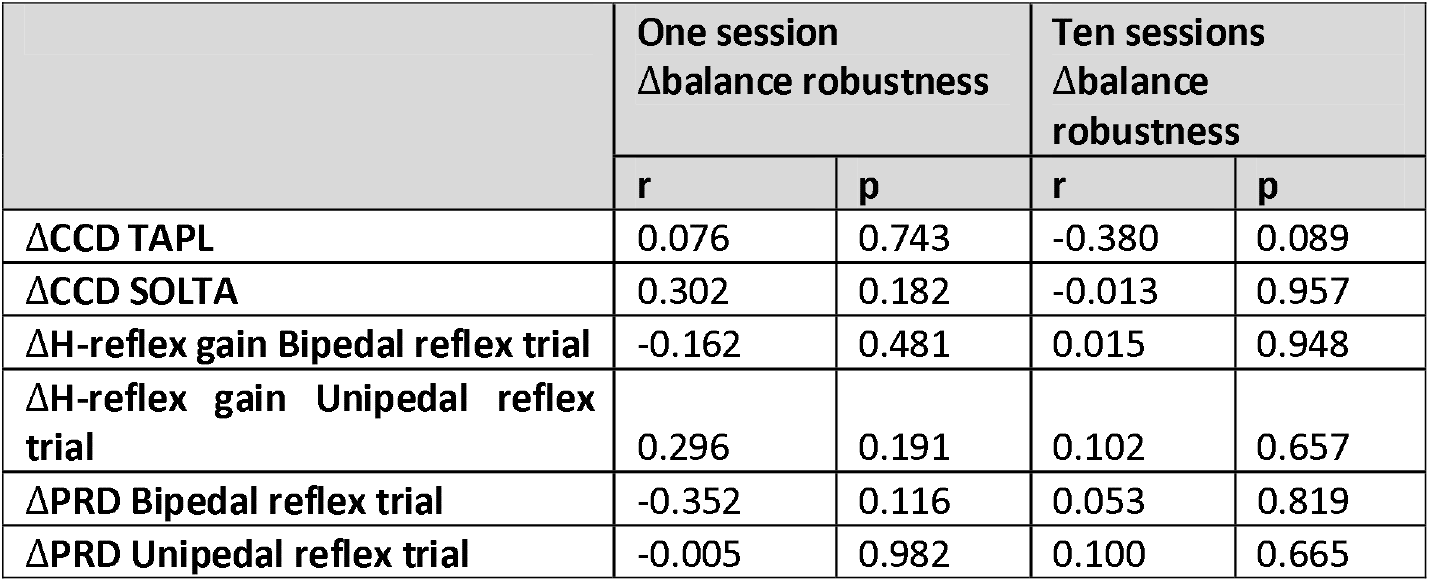
Results of the correlational analysis between the changes of co-contraction (averaged over perturbed and unperturbed trials), changes of reflexes in bipedal and unipedal reflex trials with changes of balance robustness (duration) after one session and ten sessions of training.

#### 3.3.4 Associations of balance performance with co-contraction and reflexes

All correlation results are shown in Tables 4 and 5. For co-contraction duration, at time-point Pre and Post2, SOL/TA co-contraction duration was negatively correlated with vCoM in perturbed standing (r = −0.441, p = 0.046; r = −0.471, p = 0.032, respectively), and at time-point Pre, TA/PL co-contraction duration was negatively correlated with vCoM in unperturbed standing (r = −0.453, p = 0.040). Negative correlations indicate that higher duration of co-contraction was associated with better performance (lower sway velocity). No correlations were observed between changes after one session or ten sessions of training (Table 5).

**Table 4:** Results of the correlational analysis between co-contraction with vCoM in perturbed and unperturbed, and between reflexes in bipedal and unipedal reflex trials with vCoM in bipedal and unipedal reflex trials at each Time-points of Pre, Post1, and Post2. Bold numbers indicate a significant effect.

|  | Pre vCoM |  | Post1 vCoM |  | Post2 vCoM |  |
| --- | --- | --- | --- | --- | --- | --- |
|  | r | p | r | p | r | p |
| **Perturbed** |  |  |  |  |  |  |
| CCD TAPL | -0.306 | 0.176 | -0.426 | 0.055 | 0.040 | 0.863 |
| CCD SOLTA | **-0.441** | **0.046** | -0.340 | 0.131 | **-0.471** | **0.032** |
| **Unperturbed** |  |  |  |  |  |  |
| CCD TAPL | **-0.453** | **0.040** | -0.274 | 0.228 | 0.168 | 0.462 |
| CCD SOLTA | -0.268 | 0.237 | -0.285 | 0.208 | -0.189 | 0.408 |
| **Bipedal reflex trial** |  |  |  |  |  |  |
| H-reflex gain Bipedal reflex trial | 0.066 | 0.775 | -0.137 | 0.550 | 0.310 | 0.170 |
| PRD Bipedal reflex trial | -0.375 | 0.094 | **0.583** | **0.006** | -0.275 | 0.226 |
| **Unipedal reflex trial** |  |  |  |  |  |  |
| H-reflex gain Unipedal reflex trial | 0.284 | 0.210 | -0.009 | 0.970 | 0.185 | 0.418 |
| PRD Unipedal reflex trial | -0.045 | 0.845 | 0.305 | 0.178 | -0.088 | 0.702 |

**Table 5:** Results of the correlational analysis between changes of co-contraction with changes of vCoM in perturbed and unperturbed, and changes of reflexes in bipedal and unipedal reflex trials with changes of vCoM in bipedal and unipedal reflex trials after one session and ten sessions of training

| | One session ΔvCoM | | Ten sessions ΔvCoM | |
| --- | --- | --- | --- | --- |
|  | r | p | r | p |
| **Perturbed** |  |  |  |  |
| ΔCCD TAPL | -0.089 | 0.698 | -0.258 | 0.256 |
| ΔCCD SOLTA | -0.003 | 0.988 | 0.001 | 0.997 |
| **Unperturbed** |  |  |  |  |
| ΔCCD TAPL | -0.002 | 0.993 | 0.022 | 0.925 |
| ΔCCD SOLTA | 0.306 | 0.176 | 0.085 | 0.711 |
| **Bipedal reflex trial** |  |  |  |  |
| ΔH-reflex gain Bipedal reflex trial | -0.367 | 0.101 | 0.347 | 0.124 |
| ΔPRD Bipedal reflex trial | -0.115 | 0.616 | -0.386 | 0.085 |
| **Unipedal reflex trial** |  |  |  |  |
| ΔH-reflex gain Unipedal reflex trial | -0.414 | 0.063 | 0.234 | 0.306 |
| ΔPRD Unipedal reflex trial | 0.144 | 0.531 | -0.039 | 0.868 |

For reflexes, at time-point Post1, paired reflex depression was positively correlated with vCoM in bipedal reflex trial (r = 0.583, p = 0.006), indicating that stronger paired reflex depression was associated with better performance. No correlations were observed between changes after one session or ten sessions of training (Table 5).

## 4. Discussion

We investigated the effects of one session and ten sessions of balance training in older adults on balance robustness and performance and the neural mechanisms associated with training effects. We found that only one session of balance training increased older adults’ balance robustness. Extra training sessions did not further improve but maintained the acquired robustness. In addition, balance performance in perturbed unipedal balancing was improved after only one training session, again with no further improvement over subsequent training sessions. Performance in unperturbed unipedal balancing, significantly improved over a ten sessions training period, in line with previous studies (Muehlbauer et al., 2015).

We suggest that robustness and perturbed performance outcomes are mainly limited by the ability to deal with near balance loss, while the unperturbed balance test reflects the ability to minimize sway in a situation where balance loss is not likely to occur. The fast changes in the ability to recover balance would be in line with results on perturbation training (Liu et al., 2017; Rieger et al., 2020). Improved balance performance and increased balance robustness both indicate improved balance control, however they show different aspects of balance control. To verify the latter, we calculated correlations between the outcomes of the most similar tests, the unperturbed unipedal stance and robustness tests. At pre and post2 there were no correlations between robustness and performance (Spearman’s r = − 0.285, p = 0.211; Spearman’s r = − 0.204, p = 0.375, respectively). At post1, better performance coincided with a higher robustness (Spearman’s r = − 0.759, p <0.001). This shows that the robustness measure is only partially correlated to performance. The robustness trials aimed to assess the maximal capacity to maintain balance. While it was not the intention to develop an ecologically valid test, it may reflect the ability to deal with surface instability, such as when stepping into sand, mud, or onto a carpet. We chose unipedal stance to make the test challenging enough and because balance in the unipedal stance phase of gait is crucial in daily life. Earlier studies have not used tests of robustness so it is unclear whether changes in robustness would be reflected in functional tasks. Again, they need not be as these tasks are usually submaximal and an association with performance measures is theoretically more plausible. We would expect robustness trials to be associated with the ability to prevent a fall after a major perturbation, and hence may be predictive of fall risk in daily life, but this remains to be shown.

In terms of challenge, the perturbed balance performance test and the unipedal reflex trial can be considered intermediate to the unperturbed balance test and the test for robustness. In both these tests vCoM was higher than in unperturbed unipedal and bipedal reflex trial (see supplementary material 3). During less challenging tests, such as the bipedal reflex trial and the unperturbed unipedal trial, we found no effects on balance performance after a single training session, whereas balance performance during the more challenging perturbed unipedal trial, unipedal reflex trial, and robustness trial was improved after a single session. After ten training sessions, the effects were also visible in the unperturbed unipedal trial, but not in the bipedal reflex trial.

Overall, our results suggests that balance training can increase robustness rapidly, while more sessions are needed to refine balance performance and may also be needed to maintain the acquired balance robustness and performance. Given the functional relevance of balance robustness, this finding would put into question the predominant use of balance performance in conditions with a low challenge as outcome measures of training. We note here that balance performance during bipedal reflex trial was not affected by training.

Contrary to our hypothesis, co-contraction duration of antagonistic muscle pairs was not decreased after balance training; one session of training did not change the co-contraction duration, and ten sessions of training even led to an increased co-contraction duration of SOL/TA. Moreover, cross-sectional correlation analysis showed that higher co-contraction duration was correlated with higher balance robustness and performance, primarily for the SOL/TA pair. Older adults show higher co-contraction (magnitude and duration) than young adults (Nagai et al., 2011). This increased co-contraction may be an adaptation, and training may have increased the use of this adaptation.

Higher co-contraction duration of antagonistic muscles has been shown to increase joint stiffness and serve as a zero-delay corrective response to unexpected disturbances in challenging motor tasks (Chambers & Cham, 2007). In addition, antagonistic co-contraction may reduce electromechanical delays (Oomen et al., 2015) and may improve feedback response by allowing dual control of agonist and antagonistic muscles (Saliba et al., 2020). Therefore, older adults may increase antagonistic co-contraction duration to enhance balance control. However, longitudinal analysis did not show any correlation between the changes in co-contraction duration and changes in balance robustness or performance.

Therefore, it seems that increased co-contraction duration is not the exclusive mechanism underlying improved balance after one session or ten sessions of training.

Also in contrast with our hypothesis, neither one session, nor ten sessions of training affected H-reflex gains or paired reflex depression in bipedal and unipedal reflex trials. In line with previous studies (Alizadehsaravi et al., 2020; Kim et al., 2013; Pinar et al., 2010), H-reflex gains decreased when going from bipedal to unipedal stance. This has been suggested to help in dealing with the higher postural demand of unipedal stance (Kim et al., 2012), where monosynaptic stretch reflexes may fail to contribute to maintenance of balance. However, we found stronger paired reflex depression in bipedal than unipedal stance. It has been suggested that the inhibitory effect of the first H-reflex stimulus is less when more background afferent discharge is present, which could explain the difference between unipedal and bipedal stance (Trimble et al., 2000). Alternatively, paired reflex depression may be affected by descending pathways projecting onto spinal interneurons, resulting in a larger second H-reflex (less depression) in unipedal compared to bipedal stance (Cardona & Rudomin, 1983). Functionally this decreased depression could act to facilitate responses to external perturbation, but this would be at odds with the decreased gain of the first H-reflex. Cross-sectional analyses showed that, in unipedal stance, smaller H-reflex gains were correlated with higher balance robustness, and lower paired reflex depression in bipedal stance correlated with better balance performance. Longitudinal correlational analyses did not show any significant correlation between the neuromuscular mechanisms and balance performance or the robustness. All in all, these data support that lower excitability in response to type 1a afference and stronger suppression of responses to such input is beneficial for balance control, in line with outcomes of studies in middle-aged adults (Guan & Koceja, 2011), but changes in H-reflex sensitivity or depression do not appear to account for the effect of training.

### 4.1 Limitations

Since multiple randomized controlled trials have shown the efficacy of balance training in older adults, the present study was done without a control group (Muehlbauer et al., 2015). This implies however, that we cannot exclude that some of our findings were due to repeated testing, which in itself could be seen as a form of training. However, to assess if the fast improvement is due to habituation to the tests or the training, we checked the improvement between the 3 trials at the pre-measurement time point. The results showed no significant changes between first, second and third trials at the pre-measurement time-point. This confirms that effects are a result of training (F_2,44_ =0.09, p = 0.914). Also, the finding that balance robustness did not drop two weeks after the last training session and hence five weeks after initial testing indicates that the improvement was a result of learning. Second, our hypothesis that co-contraction duration will decrease after the balance training in older adults, was based on the findings from our previous study, where we found higher co-contraction duration in older adults compared to younger adults. Hence, we used the method presented in the current study, comparable with our first study (Alizadehsaravi et al., 2020), which takes the duration of co-contraction as a percentage of when muscles are active, as determined from a reference activation. However, EMG activity of the ankle muscles during unipedal stance on a rigid surface could be reduced by training, and the higher co-contraction duration after the ten sessions of training could then be due to a lower baseline measurement at this time point. Note that co-contraction duration was measured and not co-contraction magnitude, because measurement on different days required EMG electrode replacement, which would preclude comparison of non-normalized EMG signals. Normalization to MVC was not performed as this was considered too time-consuming and potentially fatiguing.

Third, for reflex measurements it is generally recommended to elicit H-reflex between 15-40% of M_max_ (Crone et al., 1990; Knikou, 2008), while we elicited H-reflex at H_max_, in line with our previous study. However, for 20 out of 22 participants H_max_ was less than 40% of M_max_ (see supplementary material 4). Lastly, we calculated a large amount of correlations, and did not apply a correction for multiple testing while doing so. Note that there were only two participants with H_max_ higher than 40% of M_max_, therefore, we did not expect the correction to significantly change the result. Nevertheless, our results should be considered as explorative, and future, confirmative and larger preregistered studies should be undertaken to confirm our findings.

## 5. Conclusion

Previous studies showed improved balance performance as a result of balance training in both young and older adults (Cadore et al., 2013; Muehlbauer et al., 2015; van Dieën et al., 2015). In young adults improved balance control has been shown to be accompanied with decreased H-reflexes (Keller et al., 2012) and decreased co-contraction (Schinkel-Ivy & Duncan, 2018). In older adults, the mechanisms underlying improvements in balance performance and robustness after the training remain unclear. We found that one training session improved balance robustness in older adults with no further improvement after ten sessions, but the additional sessions may have contributed to the retention of improved balance robustness. Balance performance was found to be consistently improved after ten training sessions. While co-contraction duration of SOL/TA and TA/PL muscle pairs was correlated to balance performance cross-sectionally, the neural mechanisms underlying balance improvement after one or ten training sessions were not the ones studied here (i.e., co-contraction duration, H-reflex gain and peripherally induced inhibition measured with paired reflex depression).

- Leila Alizadehsaravi contributed to conceptualization, methodology, H-reflex measurement, investigation, software, formal analysis, visualization, writing original draft, review and editing, and project administration.
- Ruud A.J. Koster contributed to the investigation, co-writing the methods section of original draft, and review and editing.
- Wouter Muijres contributed to the investigation, supervision of balance training, and review.
- Huub Maas contributed to conceptualization, methodology, and review.
- Sjoerd M. Bruin contributed to conceptualization, methodology, software, review and editing, and supervision.
- Jaap H. van Dieën contributed to conceptualization, methodology, review and editing, supervision, project administration, resources and funding acquisition.

## Acknowledgments

This project has received funding from the European Union’s Horizon 2020 research and innovation programme under the Marie Skłodowska-Curie grant agreement No 721577. The research team would like to thank the individuals who participated in the experiment for the purposes of this research. SMB was funded by a VIDI grant (016.Vidi.178.014) from the Dutch Organization for Scientific Research (NWO). RAJK was supported by the European Research Council (ERC) under the European Union’s Horizon 2020 Research and Innovation Program (grant agreement number 715945 Learn2Walk). A part of this study has been presented at American society of biomechanics 2020 congress by the first author.

